# Structural adaptation of oxygen tolerance in 4-hydroxybutyrl-CoA dehydratase, a key enzyme of archaeal carbon fixation

**DOI:** 10.1101/2020.02.05.935528

**Authors:** Hasan DeMirci, Bradley B. Tolar, Tzanko Doukov, Aldis Petriceks, Akshaye Pal, Yasuo Yoshikuni, Aharon Gomez, David A. Saez, Esteban Vöhringer-Martinez, Thomas Schwander, Tobias J. Erb, Christopher A. Francis, Soichi Wakatsuki

## Abstract

Autotrophic microorganisms that convert inorganic carbon into organic matter were key players in the evolution of life on Earth. As the early atmosphere became oxygenated, microorganisms needed to develop mechanisms for oxygen protection, especially those relying on enzymes containing oxygen-sensitive metal clusters (e.g., Fe-S). Here we investigated how 4-hydroxybutyryl-CoA dehydratase (4HBD) - the key enzyme of the 3-hydroxypropionate/4-hydroxybutyrate (HP/HB) cycle for CO_2_-fixation - adapted as conditions shifted from anoxic to oxic. 4HBD is found in both anaerobic bacteria and aerobic ammonia-oxidizing archaea (AOA). The oxygen-sensitive bacterial 4HBD and oxygen-tolerant archaeal 4HBD share 59 % amino acid identity. To examine the structural basis of oxygen tolerance in archaeal 4HBD, we determined the atomic resolution structure of the enzyme. Two tunnels providing access to the canonical [4Fe-4S] cluster in oxygen-sensitive bacterial 4HBD were closed with four conserved mutations found in all aerobic AOA and other archaea. Further biochemical experiments and molecular dynamics simulations support our findings that restricting access to the active site is the key to oxygen tolerance, explaining how active site evolution drove a major evolutionary transition.

**Significance statement:** Autotrophy (primary production) was the first life strategy on Earth. Before photosynthesis (using solar energy to fix carbon dioxide), life relied on chemical reactions for energy. These chemosynthetic reactions are present in all domains of life, including archaea possessing the most energy-efficient carbon fixation pathway - the 3-hydroxypropionate/4-hydroxybutyrate cycle. This efficiency results from enzyme modifications, including enhanced enzyme stability and catalysis of multiple reactions. We reveal the first structure of aerobic 4-hydroxybutyryl-CoA dehydratase (4HBD) from ammonia-oxidizing archaea. These archaea are among the most abundant organisms on the planet, and their 4HBD active site evolved oxygen tolerance to support aerobic metabolism. This modification can provide further insight into enzyme evolution on early earth, as photosynthesis developed and began oxygenating the atmosphere.

## Introduction

The global carbon cycle and its biological influences have been deeply scrutinized as climate change intensifies ^1^. This cycle – perturbed by excessive anthropogenic carbon emissions – affects biomass in ecosystems, ocean acidity, global temperatures, and numerous other environmental factors ^1,2^. Therefore, understanding how nature both releases and sequesters carbon is key to addressing excess atmospheric carbon dioxide. Quantitatively, the assimilation of CO_2_ into organic materials (and biomass) is the most important biosynthetic process. To date, six major CO_2_-fixation pathways have been characterized ^3–5^. These autotrophic pathways have similarities, but vary in kinetics, thermodynamics, enzymology, and oxygen tolerance, indicating adaptations to various environments or niches. Such adaptations in CO_2-_fixation pathways impact fitness of autotrophic organisms, and therefore likely influenced the evolutionary divergence of early life ^6,7^. Chemolithoautotrophic ammonia-oxidizing archaea (AOA) from the phylum *Thaumarchaeota* fix CO_2_ using energy derived from the oxidation of ammonia - the first step in nitrification ^8^. AOA flourish in the nutrient-poor open ocean, increasing in abundance with depth particularly below the photic zone ^9^, where they play a significant role in CO_2_-fixation in the largest biome on Earth ^10,11^.

In order to thrive in oligotrophic environments, AOA require highly-efficient pathways especially those involving the initial conversion of inorganic carbon into biomass ^12^. Appropriately, the first cultured representative of *Thaumarchaeota* - *Nitrosopumilus maritimus* - fixes CO_2_ using a modified version of the 3-hydroxypropionate/4-hydroxybutyrate (HP/HB) cycle that was initially described in *Crenarcheaota* ^13^. This thaumarchaeal HP/HB cycle variant is the most energy-efficient aerobic CO_2_-fixation pathway presently known ^3,13^.

Though the benefits of a highly energy-efficient HP/HB cycle are clear, how this pathway and constituent enzymes evolved to adapt to dramatic changes in environmental conditions of early Earth and its atmosphere is much less understood. For example, one essential question is how Thaumarchaeota were able to maintain such an efficient CO_2_-fixation pathway as atmospheric oxygen concentrations increased to 20%. The answer may be found by investigating individual enzymes of the HP/HB cycle, particularly 4-hydroxybutyryl-CoA dehydratase (4HBD). 4HBD is a key enzyme in both aerobic thaumarchaeal and crenarchaeal HP/HB cycles, and is also found in anaerobic bacteria. In the anaerobic bacterium *Clostridium aminobutyricum*, 4HBD catalyzes a key step in 4-aminobutyrate fermentation - the non-redox dehydration of a 4-hydroxybutyryl-CoA thioester to crotonyl-CoA ^14^. In Archaea, this reaction functions canonically in the second half of the HP/HB cycle, where succinyl-CoA is processed into two acetyl-CoA molecules. The enzyme requires FAD as a cofactor, as well as a [4Fe-4S] cluster in the active site ^15^. To date, structural and biochemical characterization of 4HBD has primarily focused on previously-known anaerobic versions, specifically in *C. aminobutyricum* ^14–17^. More recently, a comparative functional characterization of 4HBD from anaerobic *C. aminobutyricum* and aerobic *N. maritimus* confirmed the stark difference in their oxygen sensitivity, ascribing it to a key evolutionary adaptation of AOA to an oxic environment ^13^. However, since no high-resolution structure exists for the 4HBD from any aerobic organism, let alone *Thaumarchaeota*, the mechanistic underpinnings behind such adaptations remain unknown. In this study, we present the first high-resolution crystal structure of 4HBD from a thaumarchaeote, *N. maritimus*. Structural comparison with that of substrate-free *C. aminobutyricum* 4HBD, assisted with all-atom molecular dynamics simulations in aqueous solution, as well as enzyme kinetics assays, shed light on the co-evolution of 4HBD during the transition from the anoxic habitats of anaerobic bacteria to oxygenated environments inhabited by aerobic *Thaumarchaeota*.

## Results

### Unusual oxygen tolerance of purified *N. maritimus* 4HBD

It is well known that [4Fe-4S] cluster-containing proteins are very sensitive to the presence of oxygen, which typically necessitates the anaerobic production, purification, and characterization of these proteins. It was predicted and demonstrated *in vitro* that *N. maritimus* 4HBD has an unusually high oxygen tolerance with a half-life of more than 10 hours ^13^. In our hands, purified 4HBD retained ∼25% activity after prolonged incubation for several days under aerobic conditions, which made it possible to perform all purification steps of crystallization samples in the presence of oxygen. The time it takes from purification to obtaining crystals is typically 3-4 days. Crystals retained their dark brown color for a few weeks, suggesting that [4Fe-4S] cluster of 4HBD has unusual stability under ambient atmospheric oxygen conditions.

### Overall Structure of *N. maritimus* 4HBD

In order to understand the molecular underpinnings of the catalytic mechanism of aerobic 4-Hydroxybutyryl-CoA dehydration, we determined the high-resolution *N. maritimus* 4HBD structure (encoded by Nmar_207) at 1.55 Å resolution. This structure shows an overall homotetrameric arrangement, which agrees with the previously reported structure of the oxygen-sensitive homolog from *C. aminobutyricum* (PDB ID 1U8V) (Figure 1). Each of the four monomers, referred to as A to D hereafter, contained an active site with one [4Fe-4S] cluster and one FAD coenzyme molecule. The space group was P12_1_1. Monomers ranged from 500-502 residues, with 456-459 of those residues facing the surface. Four large monomer-monomer interfaces were present in this structure (between monomers A-C, A-D, B-C, and B-D), ranging from 3104 to 3271 Å^2^, significant compared to the average surface area of isolated monomers ranging from 21,499 to 21,834 Å^2^. The C-B and D-A interfaces were near-identical to one another; as were the D-B and C-A interfaces. The former interfaces both contained 35 hydrogen bonds, and a similar number of salt bridges (16 in C-A, 18 in D-B). The latter contained 27 and 28 hydrogen bonds, respectively, and 24 and 25 salt bridges.

**Figure 1.**
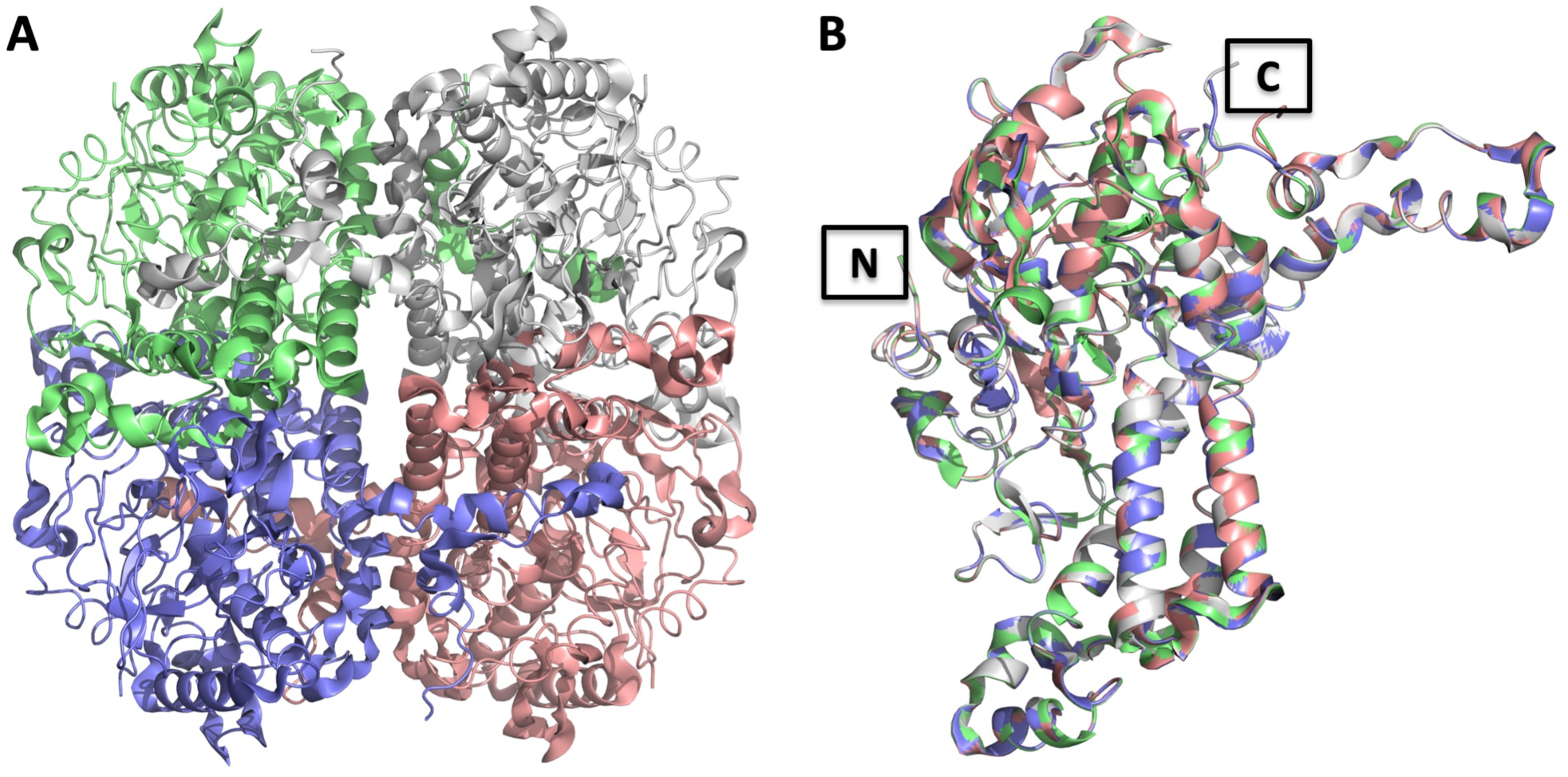
Overall structure of the *N. maritimus* 4-hydroxybutyryl-CoA dehydratase (4HBD). A) Cartoon representation of the 4HBD homotetramer. Each chain is shown in a different color. B) Superposition of all four monomers indicates the overall similarity and structural conformation of each with an overall root mean square deviation (RMSD) of 0.07-0.09 Å. The positions of the N- and C-termini of the subunits are indicated with black squares.

Within each monomer, the most prominent secondary structures are α-helices (n = 21), primarily interspersed with regions of random coil. There also existed a few regions of short β-strands, the longest of which was only 8 residues in length (Figure 2). This is similar to the *C. aminobutyricum* 4HBD structure, with the N- and C-terminal regions consisting primarily of α-helical secondary structures interspersed with random coils, and the central domain (D168-S297) containing all of the protein’s short β-strands (Figure 2).

**Figure 2.**
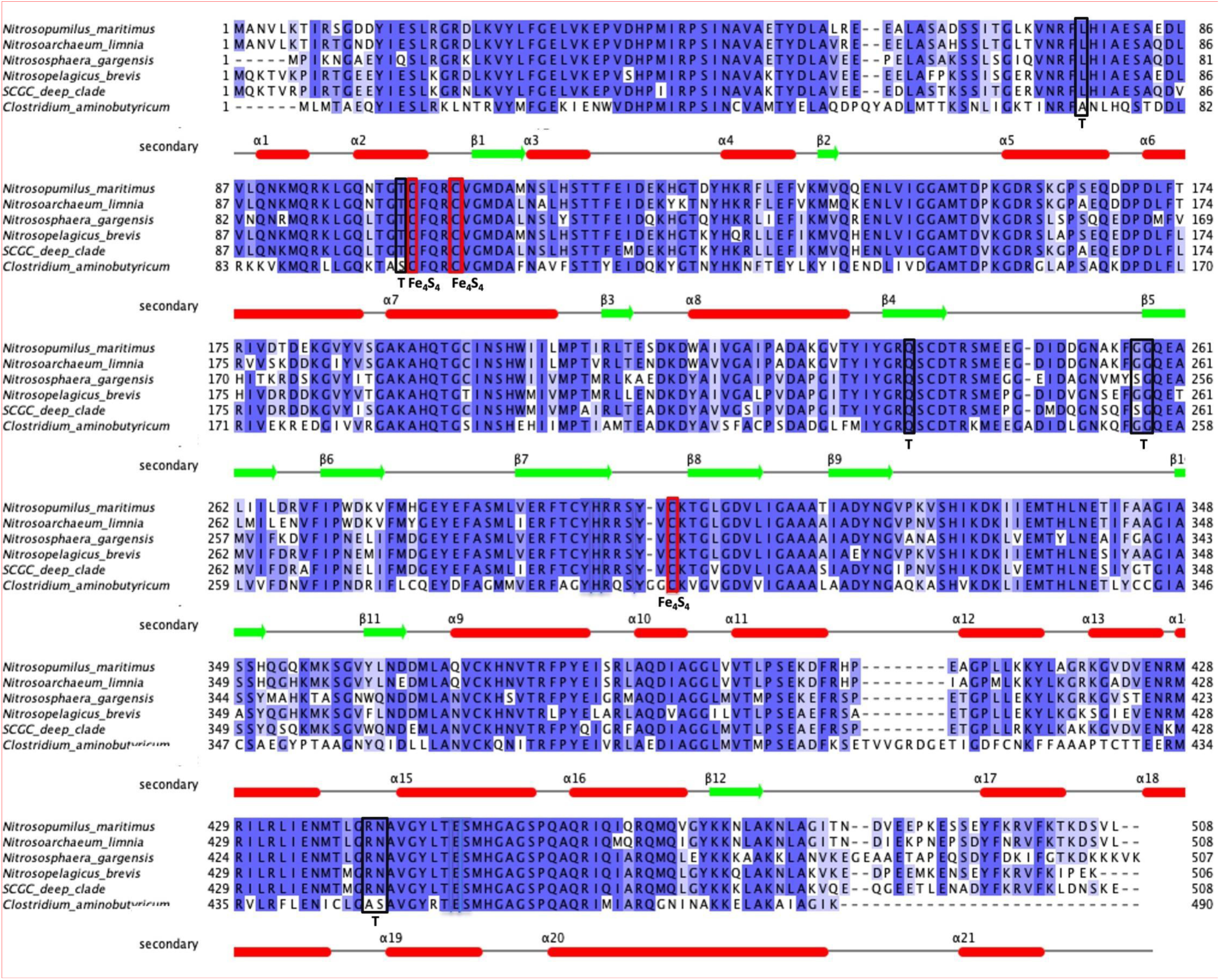
Alignment of 4-hydroxybutyryl-CoA dehydratase (4HBD) sequences from (anaerobic) *Clostridium aminobutyricum* and five representative (aerobic) ammonia-oxidizing archaea. These AOA sequences correspond to distinct ecotypes: *N. maritimus* SCM1 (marine), *Candidatus* Nitrosarchaeum limnium SFB1 (low-salinity), *Nitrososphaera gargensis* Ga9.2 (soil), *Ca*. Nitrosopelagicus brevis CN25 (oceanic water column group A, WCA), and a single-cell amplified genome ‘SCGC_deep_clade’ (oceanic water column group B, WCB). Secondary structure elements placed based on our *N. maritimus* (4HBD) structure. α-Helices are shown in red tubes and beta-strands in green arrows. Positions of important residues are indicated. The positions of catalytic cysteine residues that are involved in the [4Fe-4S] cluster coordination are enclosed by red rectangles. The positions of tunnel constriction residues T102, Q237, G257, G258, R441 and N442 on tunnel 1 and L77 on tunnel 2 are enclosed in black rectangles and indicated by letter “T” (see below and Figure 3).

The larger interfaces come together by flipping one of the two monomers from the symmetry axis by 180 degrees. In this conformation, the monomer-monomer interface is composed mainly of interactions between corresponding regions of α-helices and β-sheets.

### Constriction of the tunnels behind the [4Fe-4S] cluster protecting oxygen attack

Further inspection of the structure of the oxygen tolerant *N. maritimus* 4HBD in comparison with that of the oxygen-sensitive 4HBD from *C. aminobutyricum* revealed that both enzymes feature two tunnels that expand from the active site to the back of the protein, but also showed striking differences (Figure 3A and B). The binding pocket for the [4Fe-4S] cluster is actually part of a 45 Å long tunnel (‘tunnel 1’), which extends throughout the 4HBD monomer exiting at the other end of the protein (Figure 3B and D). Surprisingly, this tunnel is constricted severely halfway between the [4Fe-4S] and the other exit by a constellation of 6 residues, T102, Q237, G257, G258, R441, and N442 (Figure 3F). This constriction is as narrow as 2.7 Å in diameter, too small for oxygen molecules to pass. In addition to that, we observed that one water molecule at one side of the constriction makes hydrogen bonds to the carbonyl oxygen of R441, main chain nitrogen of G258, delta oxygen of N442, and epsilon oxygen of Q237. This water molecule apparently plugs the constriction site and serves as a barrier for molecular oxygen intrusion beyond this point, presumably preventing oxygen molecules from reaching the [4Fe-4S] cluster at the active site.

**Figure 3.**
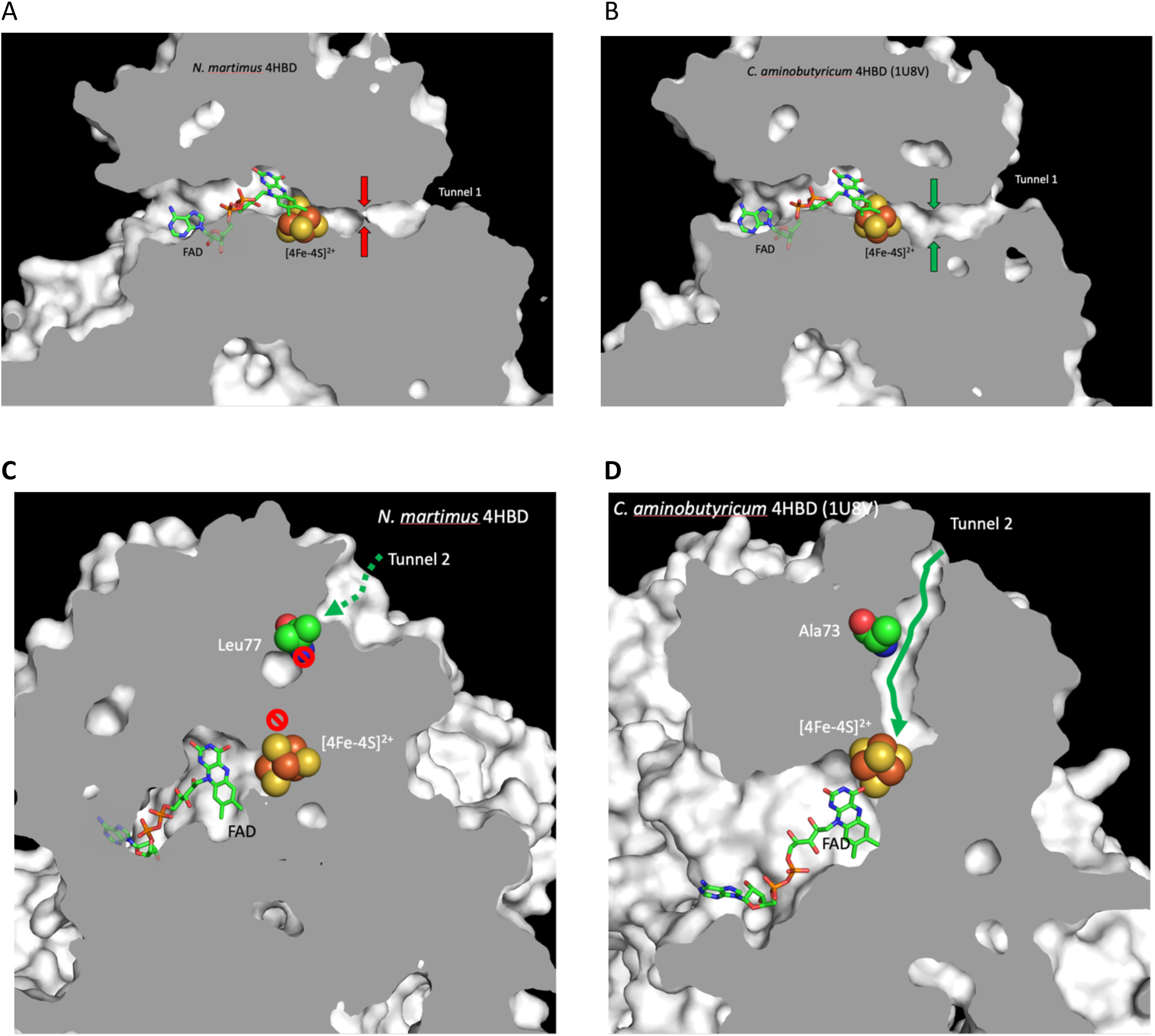
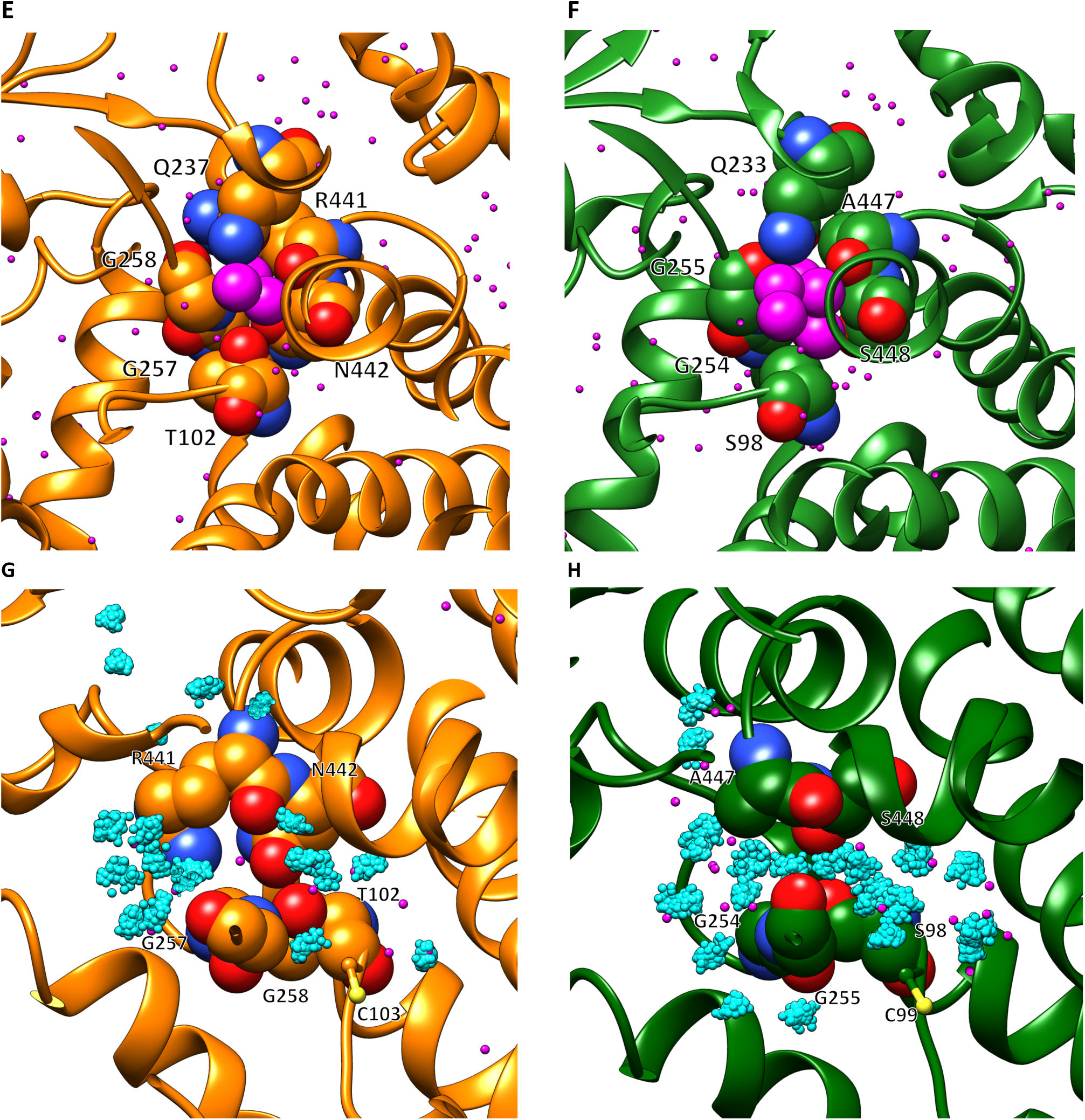
Active site tunnels reaching to the other side of the enzyme monomer likely controlling oxygen access. A) Cross section of the tunnel 1 of *N. maritimus* 4HBD including the enzyme’s active site. [4Fe-4S] cluster is shown as spheres while FAD is depicted as stick models. B) Cross section of the tunnel 1 in *C. aminobutyricum* going through the oxygen-sensitive 4HBD of active site connecting to the other side of the enzyme. The tunnel is open between the [4Fe-4S] cluster and the exit. C) Cross section of *N. maritimus* 4HBD rotated by 90 degrees around the horizontal axis of panel B, showing the tunnel 1 closed in the middle. D) Cross section viewed after a 90 degree rotation around a horizontal axis in panel B. E) 6 residues constricting the tunnel 1 at one third of the way through the tunnel, as indicated by the red vertical arrows in A. F) 6 residues forming the tunnel wall at the position, indicated by vertical green arrows in B, corresponding to the constriction site (red arrow in A) in *N. maritimus* 4HBD. G and H) Representative conformations of the closed tunnel formed by the respective residues of *N. maritimus* 4HBD and the six residues in *C. aminobutyricum* in a side view of the tunnel indicated by the vertical red arrows in A. This conformation are taken from the molecular dynamics simulations and the cyan spheres show hydration sites representing most probable positions of water molecules in the tunnel (red small spheres show the position of the crystal water molecules as reference). In E and F, other protein amino acids are shown in wire models, and water molecules as red crosses. Panels A, C, and E are drawn using molecule A of the *N. maritimus* 4HBD (this work) and B, D, and F using molecule D of *C. aminobutyricum* 4HBD (PDB 1u8v).

In contrast, the same tunnel in the oxygen-sensitive 4HBD from *C. aminobutyricum* is accessible all the way from the [4Fe-4S] cluster to the other end of the protein (Figure 3C), which is about 12 Å long and 4 to 5 Å wide. The tunnel is wide open at the position corresponding to the constriction of *N. maritimus* 4HBD. In *C. aminobutyricum* 4HBD, the cross section of the tunnel at this position is approximately 4 Å x 7 Å, surrounded by S98, Q233, G254, G255, A447, and S448. The striking differences are three residues, S98, A447 and S448 that are replaced by bulkier T102, R441 and N442, respectively, which constrict tunnel 1 in *N. maritimus* 4HBD (Figure 3E and 3F). These three residues are fully conserved in thaumarchaeal and crenarchaeal AOA (Figure 2 and Figure 4B) as well as *Plesiocystis pacifica*, which are all obligate (micro)aerobic microorganisms (Figure S4 of Berg et al. 2007)^18^. Other 4HBD sequences of anaerobic bacteria do not show strict conservation of these residues either, suggesting that 4HBD acquired oxygen tolerance by evolution of three residues to close the back entry of tunnel 1 to the active site.

**Figure 4.**
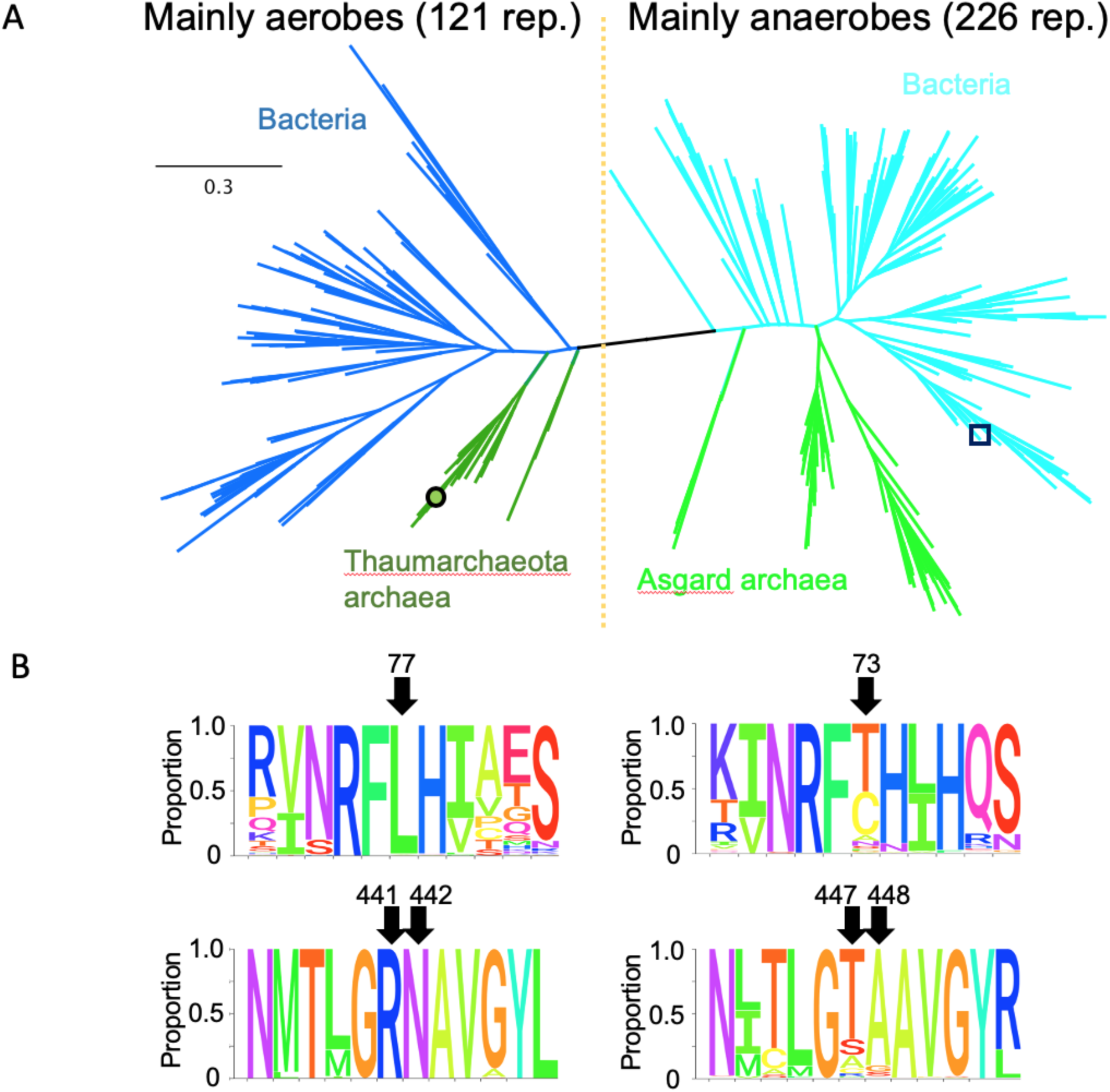
Phylogenetic analysis of the 4HBD sequences and residues constricting the back tunnels. A) Phylogenetic analysis of representative 347 4HBD homologues is shown. All sequences were first retrieved by BLASTP analysis using the Nmar 4HBD sequence as a query. We then eliminated the sequences that have length shorter than 80% of *N. maritimus* 4HBD and over 90% identity to any other sequences in the list. The 4HBD homologues derived from bacteria and archaea are shown in (dark and light) blue and (dark and light) green, respectively. The open circle and square indicate *N. maritimus* and *C. aminobutylicum* 4HB, respectively. B) The proportion of amino acid conservation for two constriction motifs corresponding to the back tunnels 1 and 2 are shown in the sequence logo at the bottom and top panels, respectively. The left panels correspond to these motifs from the clades primarily represented by aerobes. The numbers on the left panels denote the corresponding amino acid positions in the 4HBD of *N. maritimus*. The right panels correspond to these constriction motifs from the clades primarily represented by anaerobes, and the numbers denote the corresponding amino acid positions in the 4HBD of *C. aminobutyricum*.

To characterize the respective tunnels in aqueous solution we carried out molecular dynamics simulations starting from the new crystal structure of the aerobic *N. maritimus* 4HBD and anaerobic *C. aminobutyricum* 4HBD (see Methods section). After the equilibration of both proteins with their [4Fe-4S] cluster in aqueous solution similar constrictions of tunnel 1 compared to the crystal structures are observed (Figure 3F-G compared to Figure 3G-H). Hydration sites identifying the most probable water positions reveal a marked difference between the aerobic and anaerobic 4BHD (cyan spheres in Figure 3G-H). In aerobic *N. maritimus* 4HBD (Figure 3G), residue N442 and T102 disrupt a water network connecting the solvent with the [4Fe-4S] cluster observed in anaerobic 4HBD shown in Figure 3H. Water molecules inside the protein cavity close to the [4Fe-4S] cluster of aerobic 4HBD adopt similar positions as observed in the crystal structure whereas an increased number of connected hydration sites are observed in aerobic 4HBD reaching out to the solvent. We characterized the tunnel dimensions in aqueous solution with two distances involving N442, T102 and Q237 of the aerobic *N. maritimus* 4HBD and the corresponding residues in anaerobic *C. aminobutyricum* (see Supplementary Figure S3). These two distances are 1 Å shorter in aerobic 4HBD averaged over all four chains and three independent simulations providing further evidence that the tunnel is closed in aqueous solution and the [4Fe-4S] cluster not accessible from outside the protein. (see Supporting information for more details)

In *C. aminobutyricum* 4HBD, the second tunnel (‘tunnel 2’) runs perpendicular to the first, immediately behind the [4Fe-4S] cluster (Figure 3A). The corresponding tunnel of *N. maritimus* 4HBD is interrupted in two places (Figure 3B). The one interruption closer to the back exit is closed completely by the side chain of L77, which corresponds to A73 in *C. aminobutyricum* 4HBD, while the other constriction closer to the [4Fe-4S] cluster is interrupted by overall structural changes near the constriction, not by specific mutations. The branched hydrophobic side chain of L77 sticks across back tunnel 2, and is conserved throughout all aerobic AOA (Figure 2 and Figure 4B). The absence of this tunnel in the aerobic 4HBD was also confirmed by the molecular dynamics simulations of 4HBD molecule in aqueous solution, whereas the anaerobic 4BHD of *C. aminobutyricum* showed tunnels connecting the [4Fe-4S] cluster with hydration sites through this back tunnel and tunnel 1 described above (see Supporting Figure S4).

### Active Site Architecture

The crystal structure of 4HBD reveals the first structure of an oxygen-tolerant archaeal dehydratase complex, to the best of our knowledge. The active site configuration is similar to that of *C. aminobutyricum*. In both enzymes, the active site is accessed through a narrow binding channel near the surface of a monomer-monomer interface. The walls of this channel are formed by two interacting monomers. FAD rests in this channel, with its adenine ring proximal to the channel’s entry while the isoalloxazine ring is facing the [4Fe-4S] cluster. (Figure 3A, 3B). The flavin moiety is tightly constrained in the active site. For example, in one A-C active site, FAD binds to monomer A at an interface of 512 Å^2^, and monomer C at an interface of 525 Å^2^. Interestingly, this FAD was not captured in a planar geometry. Instead, we observed a bending of the isoalloxazine ring of FAD at the nitrogen N5 position, similar to the butterfly shape observed in acyl-CoA dehydrogenases suggesting a shift in the electron potential of the cofactor ^19^.

### Kinetics of active site variants

For the homologous anaerobic 4HBD from *C. aminobutyricum*, it has been reported that apart from the residues coordinating the [4Fe-4S] cluster (C99, C103, C299 and H292), four additional residues (T190, E257, Y296, and E455) are important for dehydratase activity ^17^. Therefore, we introduced mutations in the corresponding residues of the *N. maritimus* 4HBD (T194 and E260) and tested their kinetics (Table 1).

**Table 1.** Characterization of wild-type and variants of 4HBD

| 4HBD | Dehydration<br>(4-hydroxybutyryl-CoA) |  |  | Isomerization<br>(vinylacetyl-CoA) |  |  | Oligomeric<br>state |
| --- | --- | --- | --- | --- | --- | --- | --- |
|  | V<sub>max</sub><br>(U/mg) | K<sub>M</sub><br>(μM) | Relative<br>activity after<br>24h air (%) | V<sub>max</sub><br>(U/mg) | K<sub>M</sub><br>(μM) | Relative<br>activity after<br>24h air (%) |  |
| wt | 20.4 ± 0.5 | 6.4 ± 0.8 | 91 ± 9 | 108 ± 2 | 318 ± 17 | 94 ± 4 | tetrameric |
| E260Q | 0.85 ± 0.03 | 14.7 ± 1.8 | 92 ± 6 | 25 ± 3 | 318 ± 83 | 78 ± 6 | tetrameric |
| T194V | not<br>detected | n.d. | n.d. | 10 ± 1 | 284 ± 26 | 116 ± 8 | tetrameric<br>/dimeric |

n.d. = not determined. \* results the WT were obtained from three individual enzyme preparations while those of the mutants were obtained from three measurements on single preparations.

Furthermore, E260 and T194 are proposed to form a diad involved in the protonation of the FAD semiquinone. Mutation of these residues to a glutamine (E260Q) or valine (T194V), respectively, significantly reduced the dehydration activity. The T194V variant was especially affected, since T194 is able to form a hydrogen bond with the N5 of the isoalloxazine ring.

## Discussion

### Comparison of the substrate binding pockets with the bacterial counterpart

The overall structures of (aerobic) archaeal 4HBD from *N. maritimus* and the (anaerobic) bacterial homolog from *C. aminobutyricum* show a very similar architecture with RMSD of 0.54 Å between 14,738 atoms with minor differences (Supplementary Figure S1A). The *N. maritimus* 4HBD sequence has an additional 6 N-terminal residues and 24 additional C-terminal residues, while *C. aminobutyricum* 4HBD has 8 residues inserted between helices α11 and α12. These N- and C-terminal extensions of the *N. maritimus* enzyme neither affect the oligomeric state nor the overall conformational state of the 4HBD. The active sites of the two homologous enzymes, despite their differences for oxygen tolerance, display a striking similarity. Position of the critical catalytic residues E449, Y299, H295, FAD and the [4Fe-4S] cluster are highly conserved (Figure 2). In addition to the ∼58% sequence identity, the two 4HBD enzymes also share a remarkably similar surface charge distribution (Supplementary Figure S1 C,D).

### Oxygen tolerance acquired by blocking tunnels in proteins

It has been reported that blocking a protein tunnel is essential for protection in a pair of oxygen-tolerant and -sensitive [NiFe]-hydrogenases ^20–22^. The oxygen-tolerant enzyme features a tunnel that is blocked by a tryptophan at the position of the medial 3Fe-4S cluster tunnel, while the oxygen-sensitive hydrogenase has a smaller residue, phenylalanine, which does not fully close the tunnel. This situation resembles somewhat the case of the alanine to leucine mutation of tunnel 2 in 4HBD (Figure 3 A,B). In contrast, the constriction of tunnel 1 requires three mutations, S98 to T102, A447 to R441, and S448 to N442. To analyze the rigidity in the constriction of tunnel 1, we calculated the mechanical strength of the residues using the program ProPHet ^23–25^ (http://bioserv.rpbs.univ-paris-diderot.fr/services/ProPHet/). Overall, this analysis suggests that the three residues constricting tunnel 1 show above-threshold mechanical strength (Supplementary Figure S2) indicating that the constriction is rather sturdy, which corroborates well with the all-atom molecular dynamics simulations (Supplementary Figure S3).

It is not clear why 4HBDs feature such extensive tunnels at the back of their active sites containing the [4Fe-4S] clusters. Martins et al. (2004) ^15^ and Zhang et al (2015) ^17^ alluded to one of the back tunnels (tunnel 1) of *C. aminobutyricum* 4HBD as a possible escape route for the product water, but did not discuss the possible diffusion of oxygen through the two tunnels. Our structural and phylogenetic analyses of *N. maritimus* 4HBD in comparison with its oxygen-sensitive counterpart, *C. aminobutyricum* 4HBD, suggest that the 4 residues co-evolved together to close the two tunnels, probably protecting the active site of *N. maritimus* 4HBD (and other oxygen-tolerant enzymes) from an attack by oxygen.

### Concluding remarks

Although initially described in obligately anaerobic bacteria, 4HBD has been found in many aerobic Archaea ^3,13^. Though the phylogeny of crenarchaeal 4HBD used in the HP/HB cycle forms two distinct groups, all thaumarchaeal 4HBD protein sequences cluster together in a separate clade. This clade is most closely related to 4HBD from anaerobic bacteria (including *C. aminobutyricum*), indicating that evolution of the HP/HB cycle likely occurred twice in the Archaea ^13^. In contrast to anaerobic 4HBD, oxygen tolerance has been observed both with *N. maritimus* 4HBD and *Metallosphaera sedula* - a member of the *Crenarchaeota* ^13,29^. If 4HBD originated in anaerobic bacteria, then it appears that Archaea not only acquired the enzyme on two separate occasions, but also managed to convert the enzyme to a more oxygen-tolerant form twice. A structural comparison of 4HBD from *M. sedula* would allow identification of such modifications and potentially give insight for future bioengineering efforts. For now, we can observe that the kinetics of 4HBD enzyme activity (K_m_, V_max_) are nearly identical in *C. aminobutyricum* and *N. maritimus* ^13^, whereas the K_m_ of *M. sedula* 4HBD is 2-3 times higher and its V_max_ is lower by a factor of 10 ^29^.

The apparent impact of oxygen on the evolution of the HP/HB cycle is further supported by the notion that evolution of *Thaumarchaeota* coincided with the introduction of oxygen into the atmosphere and ocean in Earth’s past ^30^. Through lateral gene transfer, bacteria contributed key genes including 4HBD to the *Thaumarchaeota*. Even more interestingly, the gain of HP/HB cycle genes among AOA is distinct from non-AOA *Thaumarchaeota* and this divergence also included the gain of ammonia oxidation and cobalamin (vitamin B_12_) biosynthesis genes - hallmarks of AOA metabolism ^30^.

Within the AOA, distinct phylogenetic groupings that match environmental habitat have been described, referred to as ecotypes, for terrestrial, freshwater, and shallow and deep marine *Thaumarchaeota* ^31^. These groups are thought to have diverged from one another through a series of adaptations to their environment, although specific markers have not been identified. Some interesting insights include the gain of potassium transport for marine clades, whereas terrestrial AOA gained oxidative stress response genes ^30^. Through this diversification, key metabolic genes would also have changed, which is reflected in the enzyme kinetic differences observed between the marine (*N. maritimus*) and terrestrial ammonia monooxygenases ^12,32^. Similar changes are likely in HP/HB cycle enzymes. Therefore, by taking a structural biology approach ^33^ to investigating enzymes like 4HBD in ecologically-relevant organisms (including the five AOA ecotypes highlighted in Figure 2), we can gain a better understanding of evolution and adaptation of key enzymes in the global carbon cycle to changing environments as will be experienced with climate change.

Considering the impact of rising atmospheric CO_2_ on global climate, structural knowledge of this enzyme may prove valuable for future endeavors to address climate change by engineering natural or synthetic enzymes and pathways for the capture and conversion of CO_2_ that are more efficient than natural evolved ones ^34^. Understanding the molecular basis of oxygen tolerance in one of the key enzymes of the HP/HB cycle for CO_2_-fixation will be useful for any future engineering efforts that may form the basis for development of effective bio-based carbon sequestration technologies.

## Experimental procedures

### Cloning

The gene encoding 4-hydroxybutyryl-CoA dehydratase (4HBD; Nmar_207) was purchased from Genscript Biotech (codon-optimized with cleavable N-terminal hexa-Histidine tag). Within the gene, *Nde*I and *BamH*I endonuclease restriction sites were used to insert Nmar_207 into the pET28a vector plasmid. The plasmid was transformed into *Escherichia coli* strain BL21 (Rosetta-2), from EMD Millipore. Transformed *E. coli* were then grown overnight on agar plates (50 µg/ml kanamycin, 30 µg/ml chloramphenicol) at 37 °C.

### Protein Expression and Purification

Overnight cultures started with single colonies from agar plates in LB media supplemented with 50 µg/ml kanamycin and 30 µg/ml chloramphenicol. They were diluted 1:100 into 12 x 2 L (total 24 L) cultures for large-scale protein expression and purification. After cultures reached an optical density of 0.8 at 600 nm, IPTG (final concentration 0.7 μM) was added to induce protein expression. Cultures were incubated overnight at 16 °C, and cell pastes were collected via centrifugation for 30 mins at 4000 x g. The cells were resuspended in lysis buffer containing (50 mM TRIS HCl pH 7.0, 1 M KCl, 5 % v/v glycerol, and 0.01 % triton X100) and sonicated using a Branson Sonifier at 60% for about 20 seconds three times (Branson Ultrasonics, Danbury, CT). Following the sonication, a second centrifugation was performed by using Beckman L8M ultracentrifuge equipped with Ti45 rotor (Beckman Coulter, Palo Alto, CA) at 4 °C and 40,000 x g for 30 mins, soluble protein solution was acquired and kept at 4 °C for the rest of the purification. Protein purity and amount was estimated through SDS-PAGE.

Protein was further purified using FPLC; Ni-NTA column packed with 20 ml of Ni-NTA resin (Qiagen). The Ni-NTA column was first washed with loading buffer A (20 mM TRIS-HCl pH 7.0, 300 mM NaCl), and the bound protein was eluted in elution buffer B (500 mM imidazole, 300 mM NaCl, 50 mM TRIS, pH 7.0). Thrombin was then added to remove N-terminal cleavable hexa-histidine tag. The cleaved protein was then concentrated to 2.0 ml final concentration and loaded on the S200 column (GE healthcare). S200 size exclusion chromatography has yielded a single peak around 230 KDa, eluting at the tetrameric size 4HBD. Fractions collected, pooled and concentrated by ultrafiltration using Amicon 30 KDa size filters and quantity was measured using UV-Vis absorption spectroscopy at 280 nm. The final concentration of the protein solution was 10 mg/ml – confirmed by Nanodrop UV spectra (280 nm).

### Crystallization

For initial crystallization screens 1 μl protein solution was added to crystal screen conditions in Terasaki plates (BioExpress), in 1:1 (v/v) ratio and sitting drop geometry. The sitting drop was then covered with 20 μl of parafin oil. Initial microcrystals were obtained by extensive pre-screening with all commercially available crystallization screens at ambient temperature. Best crystals exclusively obtained by using Molecular Dimensions’ Midas screen from conditions containing 0.1 M Li_2_SO_4_, 0.1 M Tris, pH 8 with 25 % v/v Jeffamine ED-2003.

To prepare for diffraction experiments, crystals were soaked overnight in 2.5 µl of 33 % MPD as cryoprotectant (v/v) and 77 % (v/v) mother liquor. Following cryoprotection, crystals were flash-frozen by quickly plunging them in liquid nitrogen.

### X-ray Diffraction Data Collection

X-ray diffraction data for 4HBD crystals were collected on a Pilatus 6M detector at the BL12-2 beamline of the Stanford Synchrotron Radiation Light Source in Menlo Park, CA at a wavelength of 0.979 Å and -180 °C. Diffraction data for 4HBD in space group P2_1_ were collected to 1.55 Å resolution with cell dimensions a=87.5 Å b=73.0 Å c=180.5 Å α=90.0 β=98.4 γ=90.0. A single crystal was used for the 4HBD dataset. The diffraction images were processed and scaled using the *HKL2000* package ^35^. The data processing statistics are summarized in Supplementary Table S1.

### Structure Determination and Refinement

The *N. maritimus* 4HBD structure was solved by molecular replacement with the program *PHASER* ^36,37^ from the *CCP4* program suite ^38^ in space group P2_1_ to 1.55 Å resolution. The initial search model was built with the program *MODELLER* ^39^ from one of the four subunits of anaerobic *C. aminobutyricum* 4HBD (PDB ID 1u8v). After the placement of 4 of 4HBD subunits in the asymmetric unit and initial refinement with PHENIX ^40^, the model was further rebuilt with *ARP/wARP* ^41^. The resulting model was 95% complete and manually checked and completed with Coot ^42,43^. Final crystallographic refinement was performed with the program *PHENIX* ^40^. The crystallographic R_work_/R_free_ factors are 0.157/0.175 for the 4HBD dataset respectively. The stereochemical quality of the model was assessed with Procheck ^44^. The Ramachandran statistics (most favored/additionally allowed/ generously allowed/disallowed) are 92.5/7.5/0.0/0.0 %. The refinement statistics are summarized in Supplementary Table S1. Figures were generated using *PyMOL* ^45^.

### Enzyme activity assays

Dehydration activity was measured aerobically in 120 μl of 100 mM MOPS pH 7.5 buffer containing 0.2 mM NADPH, 1 μg enoyl-thioester reductase (Etr1p) and either 0.032 μg 4HBD, 15 μg E449Q, 16 μg Y299F, 3.5 μg E260Q or 15 μg T194V. The reaction was started with different concentrations of 4-hydroxybutyryl-CoA (synthesized according to the protocol by Könneke *et al* ^13^) and the consumption of NADPH was followed at 30 °C and 340 nm (Δε=6.22 mM^-1^ cm^-1^) with a Cary 60 UV-Vis spectrophotometer (Agilent Technologies, Germany).

Isomerization activity was measured aerobically in 120 μl of 100 mM MOPS pH 7.5 buffer containing 0.2 mM NADPH, 2 mM acetyl-CoA, 1 μg enoyl-thioester reductase (Etr1p), 7.6 μg 4-hydroxybutyrate CoA-transferase (AbfT) and either 0.032 μg 4HBD, 0.6 μg E449Q, 0.21 μg Y299F, 0.35 μg E260Q or 0.45 μg T194V. The reaction was started with different concentrations of 3-butenoic acid and the consumption of NADPH was followed at 30 °C and 340 nm (Δε=6.22mM^-1^ cm^-1^) with a Cary 60 UV-Vis spectrophotometer (Agilent Technologies, Germany).

To test the activity after exposure to air, the enzyme was prepared under anaerobic conditions. The initial activity was measured with the assay described above, with the substrate concentration fixed to 160 μM 4-hydroxybutyryl-CoA or 1000 μM 3-butenoic acid respectively. The enzyme stocks were split, and kept at room temperature under anaerobic or aerobic conditions for 24 h. Afterwards, activity of the anaerobic and aerobic aliquot were re-measured.

### Phylogenetic Analysis

We first performed a BLASTP analysis using the *N. maritimus* 4HBD as a query and retrieved all sequences. We then eliminated the sequences that have length shorter than 80% of *N. maritimus* 4HBD and over 90% identity to any other sequences in the list using CD-HIT (https://www.bioinformatics.org/cd-hit/)^47^ We finally performed multiple sequencing alignment and tree construction with MUSCLE and FastTree ^48^ using their default parameters. The proportions of the amino acids are plotted with R using ggseqlogo (https://omarwagih.github.io/ggseqlogo/).

### Computational Methods

Both 4HBD tetrameric crystal structure, *N. maritimus* (this work) and *C. aminobutyricum* (PDB ID: 1U8V), were solvated in an electrically neutral dodecahedral box of TIP3P water, leaving 1.5 nm between the box edge and the closest atoms of the protein. The iron-sulfur cluster and its surrounding cysteines were modeled using the parameters provided by Blachly et al. ^49^ and the rest of the protein by the amber99sb-ildn force field ^50^. Force field parameters to describe FAD were taken from the General Amber Force Field force field. The solvated system was subject to energy minimization with position restraints on the atom position of the [4Fe-4S] cluster and FAD. 100ps equilibration at 300K with the v-rescale thermostat at constant volume for the restraint system were performed followed by 100ps equilibration at 300K at constant pressure applying the Berendsen barostat at 1 bar. Finally, 300ns production dynamics at constant temperature and pressure applying the Parrinello-Rahman barostat were performed restraining the atom positions of the [4Fe-4S] cluster and FAD, correcting for any rotations and translations of the center of mass of the whole system. The Particle Mesh Ewald algorithm was used with a 1nm cut-off to treat the long range electrostatic interactions applying cubic interpolation together with a van der Waals cutoff radius of 1 nm. Bonds involving hydrogen atoms were constrained and a time step of 2 fs was used. Three independent replicas for each system were carried out and the trajectories were combined for analysis. All simulations and analysis were carried out with the GROMACS 2019 ^51^ package. Hydration site analysis was based on a 10ns NVT simulation at 300K where all protein, [4Fe-4S] cluster and FAD atoms were restrained with a 1000 kJ mol^-1^ nm^-2^ force constant in three dimensions using the SSTMap software^52^.

## Data Availability

Coordinates of the four 4HBD structure have been deposited in the Protein Data Bank under accession code 6VJR.

## Acknowledgments

H.D. acknowledges support from National Science Foundation (NSF) Science and Technology Centers grant NSF-1231306 (Biology with X-ray Lasers, BioXFEL) and Turkish Scientific and Technological Research Council grant (118C260). Use of the Stanford Synchrotron Radiation Lightsource, SLAC National Accelerator Laboratory, is supported by the U.S. Department of Energy, Office of Science, Office of Basic Energy Sciences under Contract No. DE-AC02-76SF00515. This work was in part supported by the Stanford University’s Precourt Institute for Energy, Stanford University and DOE Office of Biological and Environmental Research. The work conducted by the U.S. Department of Energy Joint Genome Institute, a DOE Office of Science User Facility, is supported under Contract No. DE-AC02-05CH11231. The SSRL Structural Molecular Biology Program is supported by the DOE Office of Biological and Environmental Research, and by the National Institutes of Health, National Institute of General Medical Sciences (including P41GM103393). The contents of this publication are solely the responsibility of the authors and do not necessarily represent the official views of NIGMS or NIH. The authors thank Mr. Yusuke Sasaki of JGI for helping with the figures. D.S., A.G. and E.V-M thank the Max Planck Society and CONICYT-PCI - MPG190003 for funding

## Author contributions

H.D., B.T., C.F., and S.W. designed and coordinated the project. H.D. prepared and characterized the samples. H.D., T.D. and S.W. analyzed data. T.S. performed kinetics experiments. H.D., T.D., A.P. helped with data collection. A.G. performed the molecular dynamics simulations and analyzed the data, D.S set up the force field of the [4Fe-4S] cluster, E.V-M. coordinated the simulations, analyzed the data and wrote parts of the manuscript. H.D., B.T., C.F., TJE, E.V-M, and S.W. wrote the manuscript with input from all of the authors.

---

**Supplementary Figure S1.**
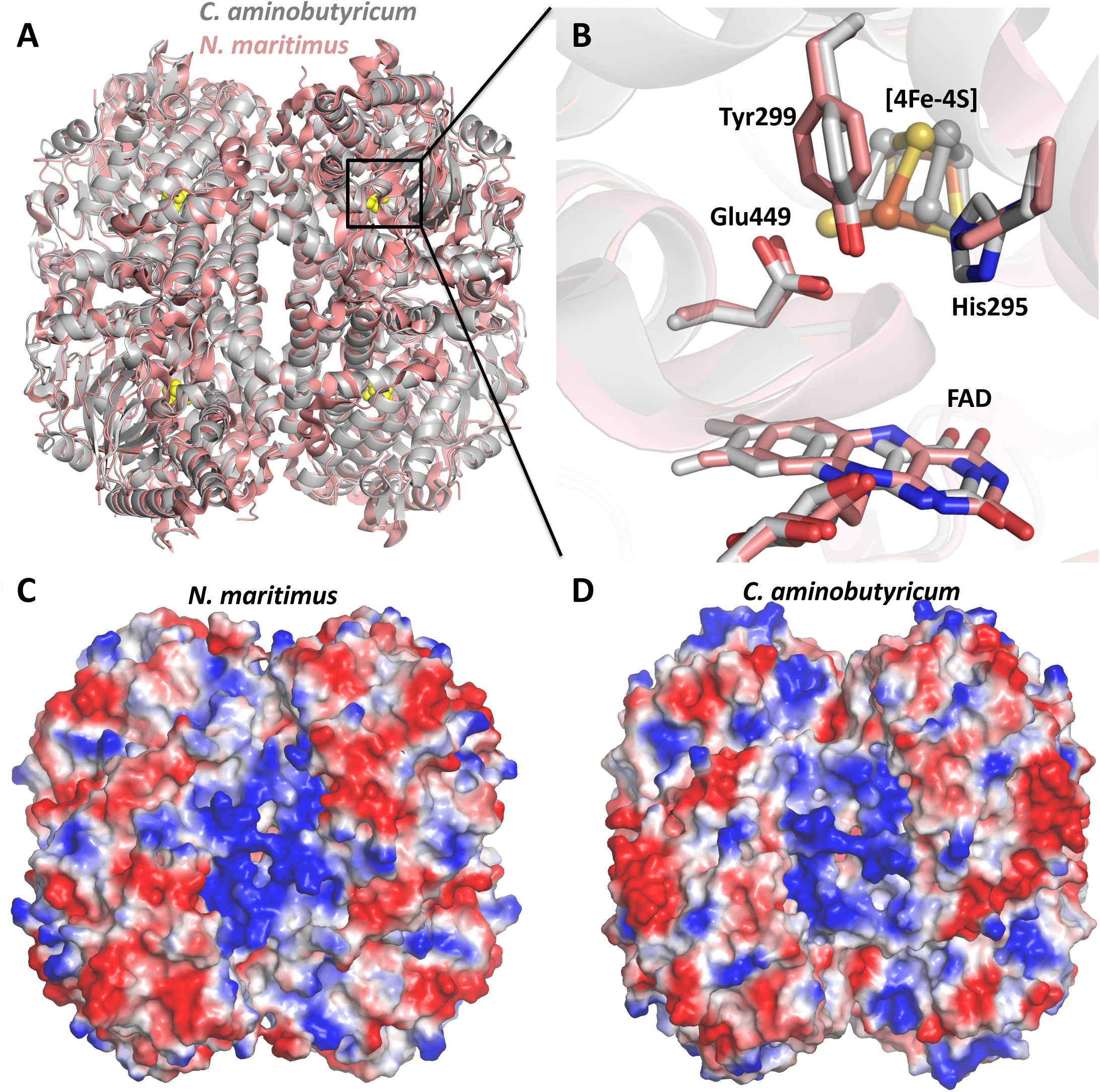
Comparison of anaerobic bacterial vs aerobic archeal 4HBD structures. A) Cartoon representation of superimposed C. *aminobutyricum* and *N. maritimus* structures, in gray and salmon colors, respectively. The overall RMSD is 0.54 Å between all the Cα residues, which indicates a high similarity between the two structures. The only notable difference is the presence of a N- and C-terminal extensions in aerobic version. B) Shows the expanded version of the inset in panel A. Positions of the active site residues E449, Y299, H295, and FAD and [4Fe-4S] cluster are highly conserved between the two 4HBD structures. C) Surface charge distribution of the *N. maritimus* 4HBD. D) Surface charge distribution of *C. aminobutyricum* 4HBD

**Supplementary Figure S2.**
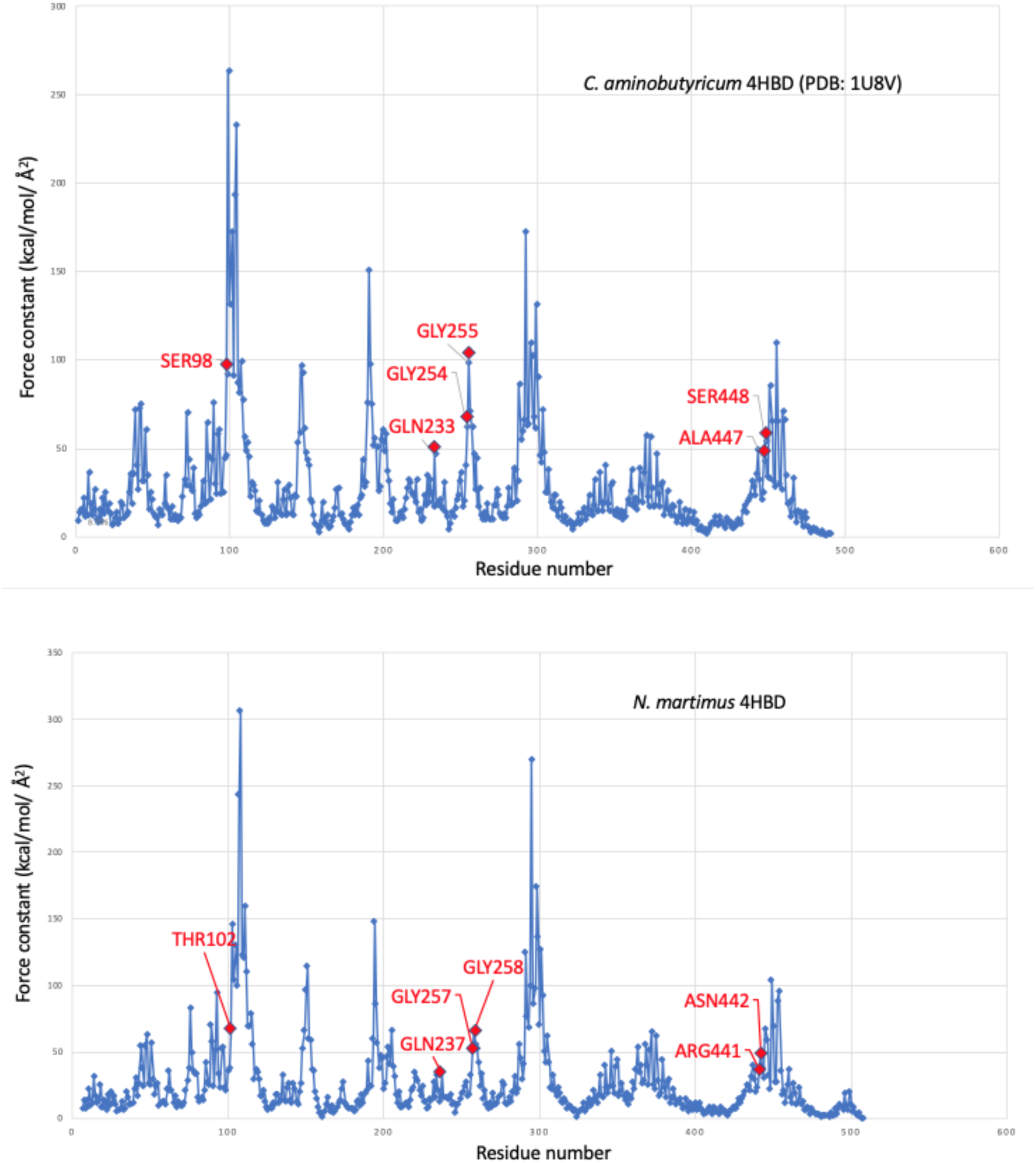
Force constants of *C. aminobutyricum (upper panel)* and *N. maritimus* (lower panel) 4HBD calculated with ProPHet (http://bioserv.rpbs.univ-paris-diderot.fr/services/ProPHet/), showing the above average force constants for the residues forming the tunnel constriction.

**Supplementary Figure 3.**
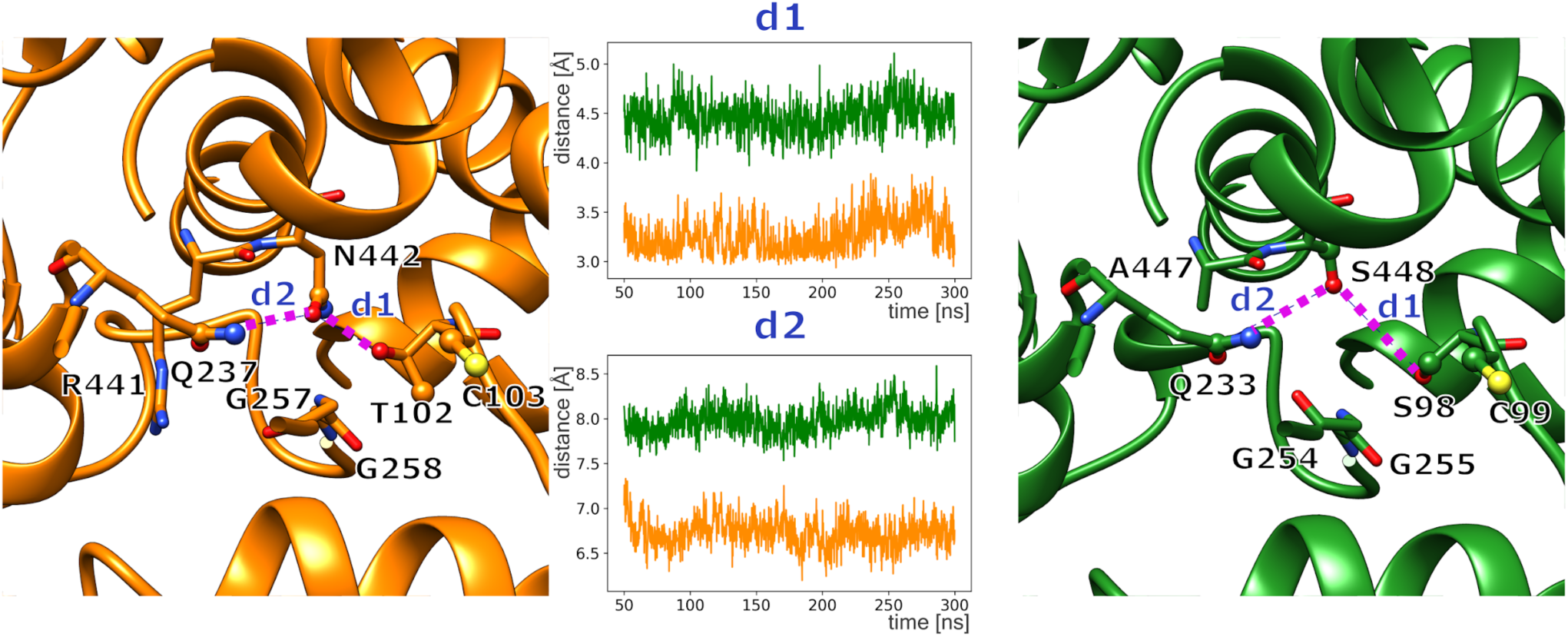
Representative conformation of the six residues forming tunnel 1 in anaerobic 4HBD from *N. maritimus* (left) and aerobic *C. aminobutyricum* (right). In the middle, the distance d1 and d2 averaged over the four monomers of the biological active tetramer as function of simulation time show larger distances and therefore an open tunnel for oxygen access in aerobic *C. aminobutyricum* (green) compared to *N. maritimus* 4HBD (orange). Distance d1 is measured between the oxygen atoms in the side chain of N442 and T102 (left) and S448 and S98 (right). Distance d2 involves the carbon atom of the amide group of Q233/Q237 and the oxygen atoms in the side chain of N442 in *N. maritimus* and S448 in *C. aminobutyricum*. Both distances characterize the dimensions of tunnel 1 to access the protein interior from aqueous solution in the molecular dynamics simulations.

**Supplementary Figure 4.**
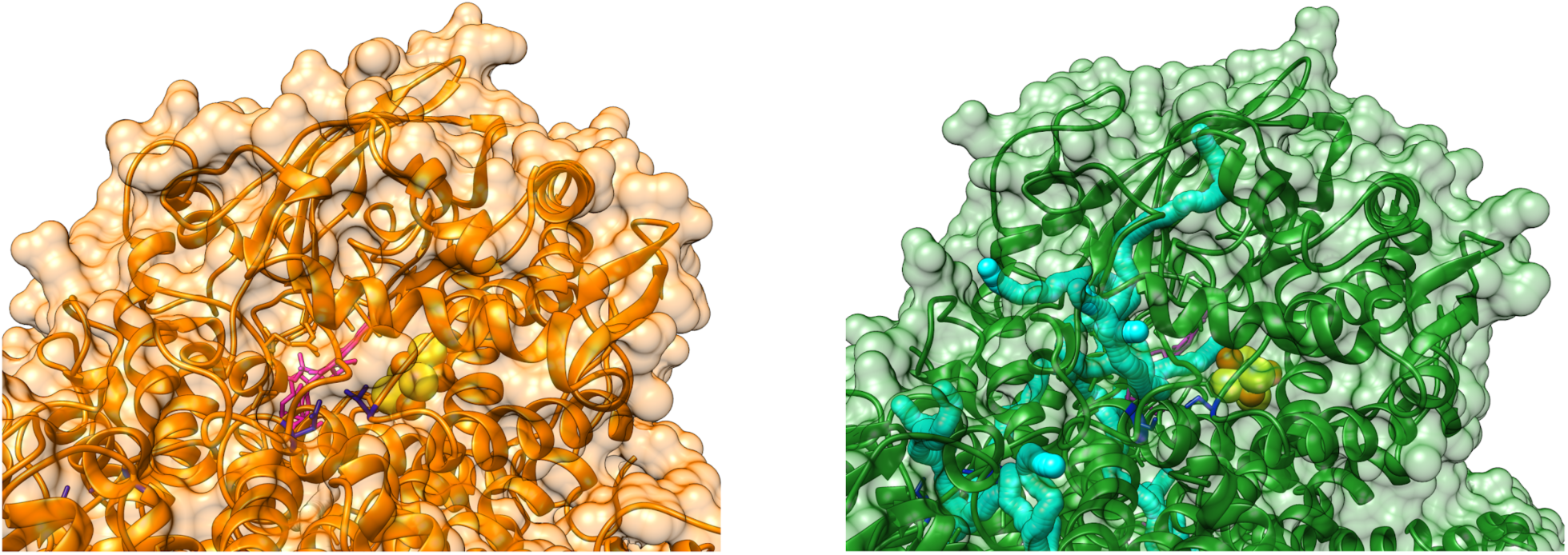
Representative conformation of one of the monomers of anaerobic 4HBD from *N. maritimus* (left) and aerobic *C. aminobutyricum* (right) from the molecular dynamics simulation. No tunnels could be found for anaerobic 4HBD (left) with the CAVER software^1^ whereas several tunnels (cyan) are observed in aerobic 4HBD (right) connecting the protein exterior with the Fe_4_S_4_ cluster (yellow and brown spheres) or FAD (magenta stick representation).

**Supplementary Table S1.**
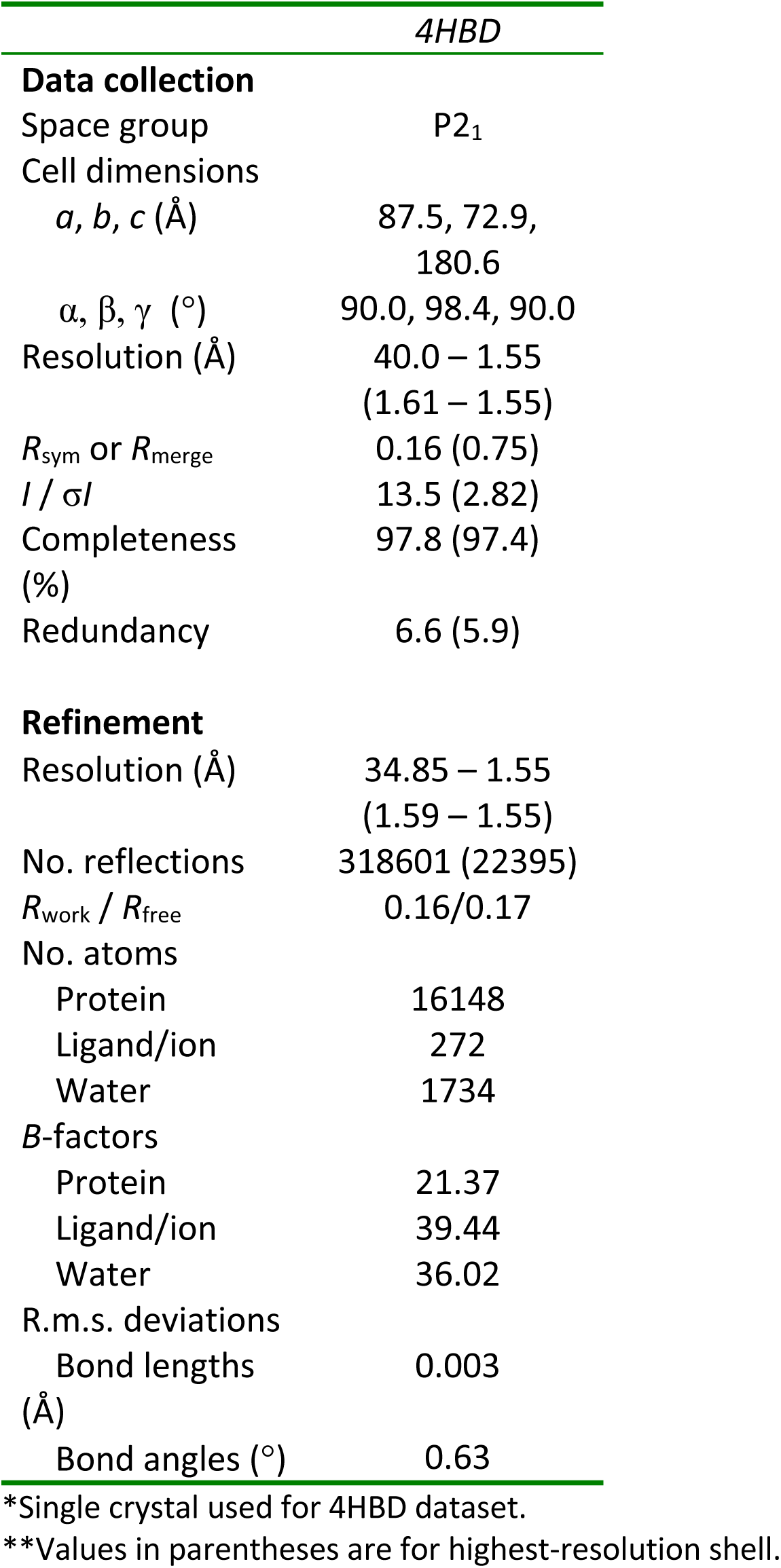
Data collection and refinement statistics

## Footnotes

1 Jurcik, A., Bednar, D., Byska, J., Marques, S. M., Furmanova, K., Daniel, L., Kokkonen, P., Brezovsky, J., Strnad, O., Stourac, J., Pavelka, A., Manak, M., Damborsky, J., Kozlikova, B. : CAVER Analyst 2.0: Analysis and Visualization of Channels and Tunnels in Protein Structures and Molecular Dynamics Trajectories., Bioinformatics, bty386, 2018., Default parameters and minimum probe radius= 1.1Å).

## References

1. Seinfeld, J. H. & Pandis, S. N. ATMOSPHERIC CHEMISTRY AND PHYSICS From Air Pollution to Climate Change SECOND EDITION. 2nd edition. Wiley, Chichester. (2006). doi: 10.1016/0016-7037

2. Liu, S. et al. Response of soil carbon dioxide fluxes, soil organic carbon and microbial biomass carbon to biochar amendment: a meta-analysis. GCB Bioenergy 8, 392–406 (2016).

3. Berg, I. A. Ecological Aspects of the Distribution of Different Autotrophic CO 2 Fixation Pathways. Appl. Environ. Microbiol. 77, 1925–1936 (2011).

4. Erb, T. J. Carboxylases in natural and synthetic microbial pathways. Applied and Environmental Microbiology (2011). doi: 10.1128/AEM.05702-11

5. Offre, P., Spang, A. & Schleper, C. Archaea in biogeochemical cycles. Annu. Rev. Microbiol. 67, 437–57 (2013).

6. Braakman, R. & Smith, E. The emergence and early evolution of biological carbon-fixation. PLoS Comput. Biol. (2012). doi: 10.1371/journal.pcbi.1002455

7. Fuchs, G. Alternative Pathways of Carbon Dioxide Fixation: Insights into the Early Evolution of Life? Annu. Rev. Microbiol. (2011). doi: 10.1146/annurev-micro-090110-102801

8. Könneke, M. et al. Isolation of an autotrophic ammonia-oxidizing marine archaeon. Nature 437, 543–546 (2005).

9. Karner, M. B., Delong, E. F. & Karl, D. M. Archaeal dominance in the mesopelagic zone of the Pacific Ocean. Nature (2001). doi: 10.1038/35054051

10. Wuchter, C. et al. Archaeal nitrification in the ocean. Proc. Natl. Acad. Sci. U. S. A. (2006). doi: 10.1073/pnas.0600756103

11. Ingalls, A. E. et al. Quantifying archaeal community autotrophy in the mesopelagic ocean usinq natural radiocarbon. Proc. Natl. Acad. Sci. U. S. A. (2006). doi: 10.1073/pnas.0510157103

12. Martens-Habbena, W., Berube, P. M., Urakawa, H., De La Torre, J. R. & Stahl, D. A. Ammonia oxidation kinetics determine niche separation of nitrifying Archaea and Bacteria. Nature (2009). doi: 10.1038/nature08465

13. Könneke, M. et al. Ammonia-oxidizing archaea use the most energy-efficient aerobic pathway for CO2 fixation. Proc. Natl. Acad. Sci. U. S. A. (2014). doi: 10.1073/pnas.1402028111

14. Scherf, U. & Buckel, W. Purification and properties of an iron-sulfur and FAD-containing 4-hydroxybutyryl-CoA dehydratase/vinylacetyl-CoA3-2-isomerase from Clostridium aminobutyricum. Eur. J. Biochem. 215, 421–429 (1993).

15. Martins, B. M., Dobbek, H., Çinkaya, I., Buckel, W. & Messerschmidt, A. Crystal structure of 4-hydroxybutyryl-CoA dehydratase: Radical catalysis involving a [4Fe-4S] cluster and flavin. Proc. Natl. Acad. Sci. U. S. A. (2004). doi: 10.1073/pnas.0403952101

16. Gerhardt, A., Çinkaya, I., Linder, D., Huisman, G. & Buckel, W. Fermentation of 4-aminobutyrate by Clostridium aminobutyricum : cloning of two genes involved in the formation and dehydration of 4-hydroxybutyryl-CoA. Arch. Microbiol. 174, 189–199 (2000).

17. Zhang, J., Friedrich, P., Pierik, A. J., Martins, B. M. & Buckel, W. Substrate-induced radical formation in 4-hydroxybutyryl coenzyme a dehydratase from Clostridium aminobutyricum. Appl. Environ. Microbiol. (2015). doi: 10.1128/AEM.03099-14

18. Berg, I. A., Kockelkorn, D., Buckel, W. & Fuchs, G. A 3-hydroxypropionate/4-hydroxybutyrate autotrophic carbon dioxide assimilation pathway in archaea. Science 318, 1782–1786 (2007). doi: 10.1126/science.1149976

19. Schwander, T., McLean, R., Zarzycki, J. & Erb, T. J. Structural basis for substrate specificity of methylsuccinyl-CoA dehydrogenase, an unusual member of the acyl-CoA dehydrogenase family. J. Biol. Chem. 293, 1702–1712 (2018).

20. Fritsch, J. et al. The crystal structure of an oxygen-tolerant hydrogenase uncovers a novel iron-sulphur centre. Nature 479, 249–252 (2011).

21. Montet, Y. et al. Gas access to the active site of Ni-Fe hydrogenases probed by X-ray crystallography and molecular dynamics. Nat. Struct. Biol. (1997). doi: 10.1038/nsb0797-523

22. Kalms, J. et al. Krypton Derivatization of an O2-Tolerant Membrane-Bound [NiFe] Hydrogenase Reveals a Hydrophobic Tunnel Network for Gas Transport. Angew. Chemie - Int. Ed. (2016). doi: 10.1002/anie.201508976

23. Lavery, R. & Sacquin-Mora, S. Protein mechanics: A route from structure to function. in Journal of Biosciences (2007). doi: 10.1007/s12038-007-0089-x

24. Sacquin-Mora, S., Laforet, É. & Lavery, R. Locating the active sites of enzymes using mechanical properties. Proteins Struct. Funct. Bioinforma. 67, 350–359 (2007).

25. Bocahut, A., Bernad, S., Sebban, P. & Sacquin-Mora, S. Frontier Residues Lining Globin Internal Cavities Present Specific Mechanical Properties. J. Am. Chem. Soc. 133, 8753–8761 (2011).

26. Darnault, C. et al. Ni-Zn-[Fe4-S4] and Ni-Ni-[Fe4-S4] clusters in closed and open α subunits of acetyl-CoA synthase/carbon monoxide dehydrogenase. Nat. Struct. Mol. Biol. 10, 271–279 (2003).

27. Doukov, T. I., Blasiak, L. C., Seravalli, J., Ragsdale, S. W. & Drennan, C. L. Xenon in and at the end of the tunnel of bifunctional carbon monoxide dehydrogenase/acetyl-CoA synthase. Biochemistry (2008). doi: 10.1021/bi702386t

28. Noor, N. D. M. et al. Redox-dependent conformational changes of a proximal [4Fe–4S] cluster in Hyb-type [NiFe]-hydrogenase to protect the active site from O 2. Chem. Commun. 54, 12385–12388 (2018).

29. Hawkins, A. B., Adams, M. W. W. & Kelly, R. M. Conversion of 4-hydroxybutyrate to acetyl coenzyme A and its anapleurosis in the Metallosphaera sedula 3-hydroxypropionate/4-hydroxybutyrate carbon fixation pathway. Appl. Environ. Microbiol. (2014). doi: 10.1128/AEM.04146-13

30. Ren, M. et al. Phylogenomics suggests oxygen availability as a driving force in Thaumarchaeota evolution. ISME J. (2019). doi: 10.1038/s41396-019-0418-8

31. Biller, S. J., Mosier, A. C., Wells, G. F. & Francis, C. A. Global Biodiversity of Aquatic Ammonia-Oxidizing Archaea is Partitioned by Habitat. Front. Microbiol. 3, (2012).

32. Dimitri Kits, K. et al. Kinetic analysis of a complete nitrifier reveals an oligotrophic lifestyle. Nature (2017). doi: 10.1038/nature23679

33. Tolar, B. B. et al. Integrated structural biology and molecular ecology of N-cycling enzymes from ammonia-oxidizing archaea. Environmental Microbiology Reports (2017). doi: 10.1111/1758-2229.12567

34. Schwander, T., Schada von Borzyskowski, L., Burgener, S., Cortina, N. S. & Erb, T. J. A synthetic pathway for the fixation of carbon dioxide in vitro. Science 354, 900–904 (2016).

35. Minor, W., Cymborowski, M. & Otwinowski, Z. Automatic system for crystallographic data collection and analysis. Acta Phys. Pol. A (2002). doi: 10.12693/APhysPolA.101.613

36. McCoy, A. J. et al. Phaser crystallographic software. J. Appl. Crystallogr. (2007). doi: 10.1107/S0021889807021206

37. McCoy, A. J. Acknowledging Errors: Advanced Molecular Replacement with Phaser. in Methods in Molecular Biology 421–453 (2017). doi: 10.1007/978-1-4939-7000-1_18

38. Winn, M. D. et al. Overview of the CCP4 suite and current developments. Acta Crystallographica Section D: Biological Crystallography (2011). doi: 10.1107/S0907444910045749

39. Webb, B. & Sali, A. Protein structure modeling with MODELLER. in Methods in Molecular Biology (2017). doi: 10.1007/978-1-4939-7231-9_4

40. Adams, P. D. et al. PHENIX : a comprehensive Python-based system for macromolecular structure solution. Acta Crystallogr. Sect. D Biol. Crystallogr. 66, 213–221 (2010).

41. Langer, G., Cohen, S. X., Lamzin, V. S. & Perrakis, A. Automated macromolecular model building for X-ray crystallography using ARP/wARP version 7. Nat. Protoc. (2008). doi: 10.1038/nprot.2008.91

42. Emsley, P., Lohkamp, B., Scott, W. G. & Cowtan, K. Features and development of Coot. Acta Crystallogr. Sect. D Biol. Crystallogr. (2010). doi: 10.1107/s0907444910007493

43. Emsley, P. & Cowtan, K. Coot: Model-building tools for molecular graphics. Acta Crystallogr. Sect. D Biol. Crystallogr. (2004). doi: 10.1107/S0907444904019158

44. Laskowski, R. A., MacArthur, M. W., Moss, D. S. & Thornton, J. M. ProCHECK: a program to check the stereochemical quality of protein structures. J. Appl. Crystallogr. (1993). doi: 10.1107/s0021889892009944

45. DeLano, W. L. References. Hypertens. Res. 37, 362–387 (2014).

46. Fish, W. W. Rapid Colorimetric Micromethod for the Quantitation of Complexed Iron in Biological Samples. Methods Enzymol. (1988). doi: 10.1016/0076-6879(88)58067-9

47. Li, W. & Godzik, A. Cd-hit: a fast program for clustering and comparing large sets of protein or nucleotide sequences. Bioinformatics 22, 1658–1659 (2006).

48. Price, M. N., Dehal, P. S. & Arkin, A. P. FastTree 2 - Approximately maximum-likelihood trees for large alignments. PLoS One (2010). doi: 10.1371/journal.pone.0009490

49. Blachly, P. G., de Oliveira, C. A. F., Williams, S. L. & McCammon, J. A. Utilizing a Dynamical Description of IspH to Aid in the Development of Novel Antimicrobial Drugs. PLoS Comput. Biol. 9, e1003395 (2013).

50. Maier, J. A. et al. ff14SB: Improving the Accuracy of Protein Side Chain and Backbone Parameters from ff99SB. J. Chem. Theory Comput. (2015). doi: 10.1021/acs.jctc.5b00255

51. Abraham, M. J. et al. Gromacs: High performance molecular simulations through multi-level parallelism from laptops to supercomputers. SoftwareX (2015). doi: 10.1016/j.softx.2015.06.001

52. Haider, K., Cruz, A., Ramsey, S., Gilson, M. K. & Kurtzman, T. Solvation Structure and Thermodynamic Mapping (SSTMap): An Open-Source, Flexible Package for the Analysis of Water in Molecular Dynamics Trajectories. J. Chem. Theory Comput. (2018). doi: 10.1021/acs.jctc.7b00592

